# Identification and Characterisation of the CD40-Ligand of *Sigmodon hispidus*

**DOI:** 10.1101/337089

**Authors:** Marsha S. Russell, Abenaya Muralidharan, Louise Larocque, Jingxin Cao, Yvon Deschambault, Jessie Varga, Sathya N. Thulasi Raman, Xuguang Li

## Abstract

Cotton rats are an important animal model to study infectious diseases. They have demonstrated higher susceptibility to a wider variety of human pathogens than other rodents and are also the animal model of choice for pre-clinical evaluations of some vaccine candidates. However, the genome of cotton rats remains to be fully sequenced, with much fewer genes cloned and characterised compared to other rodent species. Here we report the cloning and characterization of CD40 ligand, whose human and murine counterparts are known to be expressed on a range of cell types including activated T cells and B cells, dendritic cells, granulocytes, macrophages and platelets and exerts a broad array of immune responses. The cDNA for cotton rat CD40L we isolated is comprised of 1104 nucleotides with an open reading frame (ORF) of 783bp coding for a 260 amino acid protein. The recombinant cotton rat CD40L protein was recognized by an antibody against mouse CD40L. Moreover, it demonstrated functional activities on immature bone marrow dendritic cells by upregulating surface maturation markers (CD40, CD54, CD80, and CD86), and increasing IL-6 gene and protein expression. The availability of CD40L gene identity could greatly facilitate mechanistic research on pathogen-induced-immunopathogenesis and vaccine-elicited immune responses.

## 1. Introduction

The cotton rat (*Sigmodon hispidus*) was first used in polio research in the 1930s [1], and throughout the last century, it has proven to be an excellent model for biomedical research [2] [3] [4]. Historically in biomedical research, the mouse has been exploited as the default animal model. This is in part due to its well defined immunological and genetic information, cost-effectiveness, and abundant inbred strains and research reagents. However, the use of mice as models to study infectious diseases has its limitation since mice are not naturally infected by most human pathogens. On the other hand, cotton rat is susceptible to many human pathogens and is the ideal model of choice for measles (paramyxovirus) [5], herpes simplex (oral and ophthalmic) [6], influenza (orthomyxovirus) [7] [8], HIV-1 [9], RSV (respiratory syncytial virus) [10], adenovirus [11] [12], human parainfluenza [13], and human metapneumovirus [14]. This model has been valuable for adenovirus-based gene replacement therapy research [15] [16] and was also proven to be indispensable in pre-clinical evaluation of the prophylactic antibodies (RespiGam^®^ [17] and Synagis^®^ [18]. Indeed, the cotton rat model was found to be valuable in terms of its biological and immunological relevance, it was deemed unnecessary to test the adenovirus-based gene therapy and the Synagis^®^ prophylactic treatment against RSV disease in non-human primate prior to the human trials [19] [20].

A number of methods and reagents have been developed for the analysis of immune responses in cotton rats over the last decade. Up to date, more than 200 genes encoding cytokines, chemokines, cell surface markers and regulatory molecules have been cloned, with various related research reagents being commercially available. As a result, the use of cotton rats in pathogenesis studies addressing mechanistic questions has significantly increased. Nevertheless, the gene encoding CD154 and CD40 ligand (CD40L), remains elusive.

CD40L plays a critical role in orchestrating immune responses against pathogens. Depending on the post-translational modification, the murine CD40L is a 32-39 kDa type II membrane glycoprotein that was initially identified as a surface marker exclusive to activated CD4^+^ T cells [21] [22]. It is a member of the TNF superfamily consisting of a sandwiched extracellular structure composed of a β-sheet, α-helix loop, and a β-sheet, allowing for the trimerization of CD40L, an additional feature of the TNF family of ligands [23]. Since its initial discovery, CD40L has been shown to be not only expressed on CD4+ T cells, but on dendritic cells (DCs) [24], B cells [25], and platelets [26].

It has been shown that upon interacting with its receptor, CD40, CD40L induces profound effects on T cells, DCs, B cells, endothelial cells, as well as many cells of the hematopoietic and non-hematopoietic systems. Moreover, when CD40L engages CD40 on the surface of DCs, it promotes cytokine production, the induction of cell surface co-stimulatory molecules, and facilitates the cross-presentation of antigen by these cells [27], enabling DCs to mature and effectively induce the activation and differentiation of T cells. When CD40L engages CD40 on the surface of B cells, it promotes germinal center formation, immunoglobulin (Ig) isotype switching, somatic hypermutation to enhance antigen affinity, and lastly, the formation of long-lived plasma cells and memory B cells [28].Various studies have been conducted to utilize gene delivery of CD40L to DCs and tumor cells for tumor immunotherapy. It was found that expression of CD40L in a small proportion of tumor cells was sufficient to generate a long-lasting systemic anti-tumor immune response in mice that was shown to be dependent on cytotoxic T lymphocytes [29] [30].

Here we report the successful cloning of the gene encoding cotton rat CD40L (crCD40L); we also expressed and purified the CD40L produced in mammalian cells. Further characterisation of the recombinant cotton rat CD40L revealed its functional activities in promoting DC maturation and cytokine production.

## 2. Materials and Methods

### 2.1 Animals and Ethics Statement

6–7 weeks old cotton rats were obtained from an inbred colony maintained at Envigo (USA). All animal experiments were conducted in accordance with Health Canada institutional guidelines and the approval of the Animal Care and Use Committee.

### 2.2 Isolation and sequence determination of cotton rat CD40L cDNA

The spleens from three naïve cotton rats were removed aseptically and snap frozen in liquid nitrogen. The spleens were homogenized individually with a TissueLyser II (Qiagen) and total RNA extracted using the RNeasy Mini kit (Qiagen) with on-column DNase digestion according to the user’s manual. The 3’ RACE system (Life Technologies) was then used with to amplify the 3’ portion of the cotton rat CD40L from the total RNA according to the manufacturer’s instructions. A gene specific primer (5’ – GGACTCTATTATGTCTACACCCAAGTCACCTTCTG -3’) was derived from a consensus sequence aligning the rat (*Rattus norvegicus* UniProt: Q9Z2V2), mouse (*Mus musculus* UniProt: P27548), and golden hamster (*Mesocricetus auratus* XM_005084522.3). Following first strand cDNA synthesis, the 3’ portion of the cotton rat CD40L mRNA was PCR amplified using the consensus sequence derived gene specific primer and the abridged universal amplification primer with an annealing temperature at 56°C. The reverse complementary sequence of this primer was then used as a reverse primer with the forward primer (5’ - GATAGAAACATACAGCCAACCTTCTCCCAGATC -3’) to amplify the 5’ portion of the cotton rat CD40L mRNA with an annealing temperature of 57°C.

All amplified fragments were sequenced with BigDye Terminator v.3.1 Cycle Sequencing kit (ThermoFisher cat # 4336917). Briefly, samples were amplified in a PTC-200 thermal cycle (MJ Research) with the following program: 26 cycles of 1°C/S to 96°C, 96°C for 10 seconds, 1°C/S to 50°C, 50°C for 5 seconds, 1°C/S to 60°C, 60°C for 4 minutes. The samples were cleaned using DyeEx 2.0 Spin kit (Qiagen cat # 63204) and loaded onto a 3130xl Genetic Analyzer (Applied Biosystems). Raw sequencing data was edited by the instrument’s software (ThermoFisher 3130xl Genetic Analyzer Data Collection Software v3.0), and then imported into GeneCodes Sequencher v4.6.1 sequencing analysis software for further editing. The final sequenced contigs are then imported to NCBI BLAST (https://blast.ncbi.nlm.nih.gov/Blast.cgi) to confirm the identity.

### 2.3 Sequence and phylogenetic analysis

Putative conserved domains, trimer interface, and receptor binding sites were determined by performing a standard protein BLAST (blastp algorithm; https://blast.ncbi.nlm.nih.gov). The sequences producing significant alignments were imported into Geneosis software, (Auckland, New Zealand). Multiple alignment was conducted as previously described [31], with phylogenetic analysis using Geneosis Pro 5.6.7.

### 2.4 Cloning of crCD40L into Vaccina Virus expression system

Once the mRNA sequence was confirmed, a construct was designed beginning with a kozak sequence (5’-CACCGCCGCCACC – 3’), followed by a secretion signal consisting of 23 amino acid (aa) (MLLAVLYCLLWSFQTSAGHFPRA) from the human tyrosinase signal peptide as previously described [32]. This is followed by six histidine residues to facilitate protein purification. Following this sequence, a 27-aa fragment from the bacteriophage T4 fibritin trimerization motif was added [33] and finally connected to the full length 783bp open reading frame (ORF) of the cotton rat CD40L sequence at the C terminus. This construct was synthesized and cloned into pUC57 (Biobasic, Markham, ON).

Generation of a recombinant vaccinia virus expressing cotton rat CD40L protein construct was achieved using a vaccinia virus E3L and K3L double deletion mutant virus as the parental virus and taterapoxvirus K3L as the positive selection marker (Jingxin Cao, unpublished information). Briefly, the recombination plasmid vector for expression of the CD40L construct gene consists of the homologous flanking vaccinia DNA sequences targeting vaccinia A45R gene (SOD homolog); the CD40L construct gene driven by a modified vaccinia H5 promoter (Vaccine 1996, 14:1451), and taterapoxvirus 037 gene driven by vaccinia K3L promoter as the positive selection marker. The recombination vector was transfected into a HeLa PKR knockout cells infected with a vaccinia virus with both E3L and K3L genes deleted. Selection and purification of the recombinant vaccinia virus expressing the CD40L was done in BHK21 cells.

### 2.5 Western Blot

Expression of the CD40L protein was confirmed by Western blotting using His-tag Ab. Cell monolayers were lysed in sample buffer and homogenized using QIAshredder columns (Qiagen). Western blotting was performed using 4 to 15% TGX gel and Tris/Glycine/SDS running buffer (Bio-Rad Laboratories Inc.), and the protein samples were transferred to Immobilon-FL PVDF membranes (Millipore). Protein was detected with Tetra-HIS Ab (Qiagen) and goat anti-mouse IRDye-800CW (LiCor). Membranes were developed using the Odyssey system (LiCor).

### 2.6 Expression and Purification of recombinant crCD40L

The vaccinia virus carrying the crCD40L gene was propagated in BHK21 cells. The cells were collected and washed with PBS once and then lysed with a denaturing buffer (10 mM Tris– HCl, 100 mM sodium phosphate, 6 M guanidine hydrochloride, 10 mM reduced glutathione, pH 8.0) and disrupted by sonication on ice using a Branson sonifier 150 (ThermoFisher, Waltham, MA) at level 1 for two 10sec bursts with 1min rest on ice between. After separation of cell debris, the supernatant was added to a slurry of Ni-NTA resin (Qiagen, Mississauga, ON, Canada) (10 mL resin bed) and stirred at room temperature for 30 min before loading into a column. The column was purified using an AKTA purifier (Amersham Biosciences) with Unicorn 5.3 software (Amersham Biosciences). Refolding was accomplished under oxidative conditions with a gradient of denaturing buffer to buffer B (buffer B: 10 mM Tris–HCl, 100 mM sodium phosphate, pH 7.8) over 10 column volumes (CVs). The column was then washed with three CVs of buffer B + 60 mM imidazole (pH 7.8) to remove unspecific binding. The protein was eluted off the column with buffer B + 250 mM imidazole (pH 7.8). The resulting protein was dialysed against PBS pH 7.5 and then confirmed by western blot.

### 2.7 Enzyme-linked immunosorbant assay (ELISA)

96-well plates were coated with either recombinant mouse CD40L (R&D Systems) or the recombinant crCD40L protein 2ug/ml in 100µl PBS. Plates were washed with wash buffer (PBS-0.1% tween-20) and then blocked with 200µl/well blocking buffer (PBS containing 0.1% Tween 20 and 3%IgG Free BSA) for 1 hour at 37°C. Plates were washed with wash buffer and incubated at 37°C for 1 hour with 100µl/well goat α-mouseCD40L (R&D Systems) 2ug/ml in blocking buffer. Plates were subsequently washed and incubated at 37°C for 1 hour with 100µl/well with rabbit anti-goat IgG (Zymed). Plates were washed again and incubated for 10 min in the dark with 100µl/well 3,3’5,5’-tetramethylbenzidine substrate (New England Bio Labs). The reaction was stopped with Stop solution (New England Bio Labs) and absorbance was read at 450nm on a BioTek Synergy 2 plate reader.

### 2.8 Maturation and activation analysis of mouse bone marrow DC

Primary bone marrow cells from Balb/c mice (Chicago, IL) were thawed and cultured in dendritic cell medium from manufacture (Cell Biologics M7711) supplemented with GMCSF (Cell Biologics) without IL-4 at 4×10^5^ cells/well in a volume of 200µl. The cells were treated with 0.5µg/ml recombinant mouse CD40L (Preprotech, Montreal, QC) or the recombinant crCD40L protein at 0.5µg/ml, 5µg/ml, or 50µg/ml. Forty hours later, flow cytometry was performed on a BD LSRFortessa cell analyser after 2 × 10^5^ cells/tube were stained using CD11c-PE-CF594, CD54-FITC, CD40-BV786, CD80-BV421, and CD86-BV711 antibodies. All antibodies were purchased from BD Biosciences. The resulting spectra were analysed using FACSDiva version 8.0.1 software.

To assess IL-6 mRNA production of immature bone marrow murine DCs in response to targeting by recombinant crCD40L, quantitative real-time PCR was conducted on an ABI Prism 7500 Fast Sequence detection system (Applied Biosystems). TaqMan assay reagent kits (Applied Biosystems) were used that contain pre-standardized primers and TaqMan MGB probes for IL-6 and 18S which were used to normalize the data. Total RNA was isolated from 8×10^5^ stimulated bone marrow DCs using the RNeasy Mini Kit (Qiagen) according to manufactures instructions. The isolated RNA was used to make cDNA using the Superscript III First-Strand Synthesis System for RT-PCR (Invitrogen) according to manufacturer’s instructions. The cDNA was then subjected to quantitative PCR using the TaqMan Fast Advanced Master Mix (Applied Biosystems) according to manufactures instructions. Samples were run in duplicate and C_t_ values were obtained. Fold change over unstimulated DCs was calculated using the 2^-ΔΔ*CT*^ method of relative quantification [34] using 18S as the housekeeping reference gene. To investigate IL-6 secretion by murine bone marrow DCs, supernatant from forty hour stimulated cultures were collected and assayed using the Mouse IL-6 DuoSet ELISA Kit (R & D Systems) following the manufacturer’s protocol.

## 3.Results and Discussion

### 3.1 Sequence determination of the cotton rat CD40L coding sequence

The complete mRNA sequence of CD40L was obtained in two steps (Fig 1). A sequence corresponding to nucleotides 535 through to the poly-A tail was obtained using the 3’ RACE kit and mRNA as starting material, which was isolated from cotton rat splenocytes and a rodent consensus sequence as a primer. This portion of the sequence has the 3’ un-translated region of the mRNA as well as the stop codon. The 5’ end of the protein was obtained in the next step by PCR amplification of the cDNA obtained in the first step with the 3’ RACE kit and the reverse complement of the consensus sequence primer and a second consensus sequence primer designed to bind to the beginning of the CD40L mRNA. The 783bp ORF encodes 260aa followed by a stop codon.

**Figure 1:**
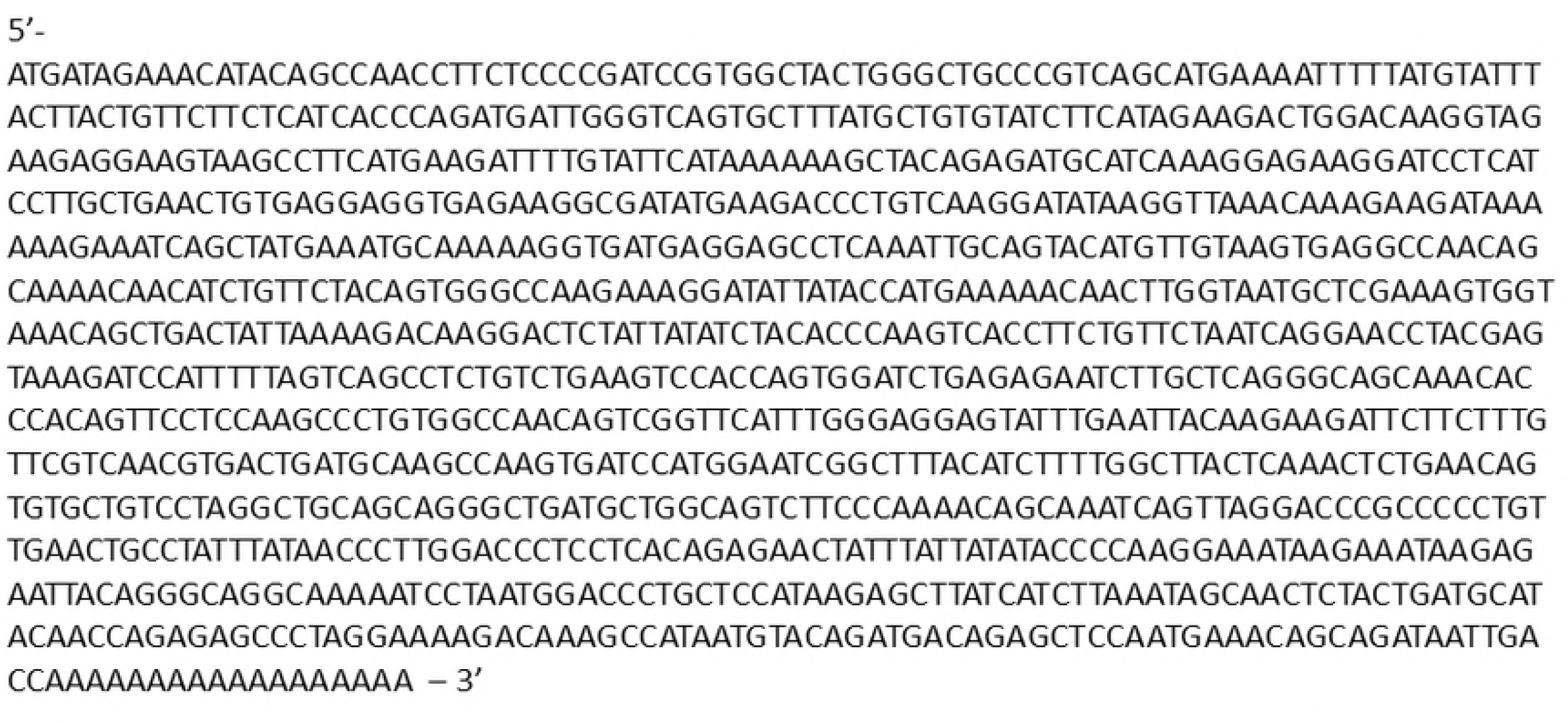
Cotton Rat CD40L mRNA sequence. The sequence was determined using 3’ RACE of mRNA extracted from the spleen of a cotton rat.

Comparison of the sequenced CD40L gene revealed that the crCD40L coding sequence shares 93%, 89%, and 83%, identity with golden hamster, rat, and mouse, respectively. At the amino acid (aa) level, the corresponding identities are 91%, 82%, and 82%, Fig 2a. At both the mRNA and aa levels, the crCD40L shared the closest similarity with Peromyscus maniculatus bairdii (or deer mouse) at 93% and 92% respectively. When sequence homology analysis is performed, crCD40L clusters with other members of the Cricetidae family Fig 2b.

**Fig 2:**
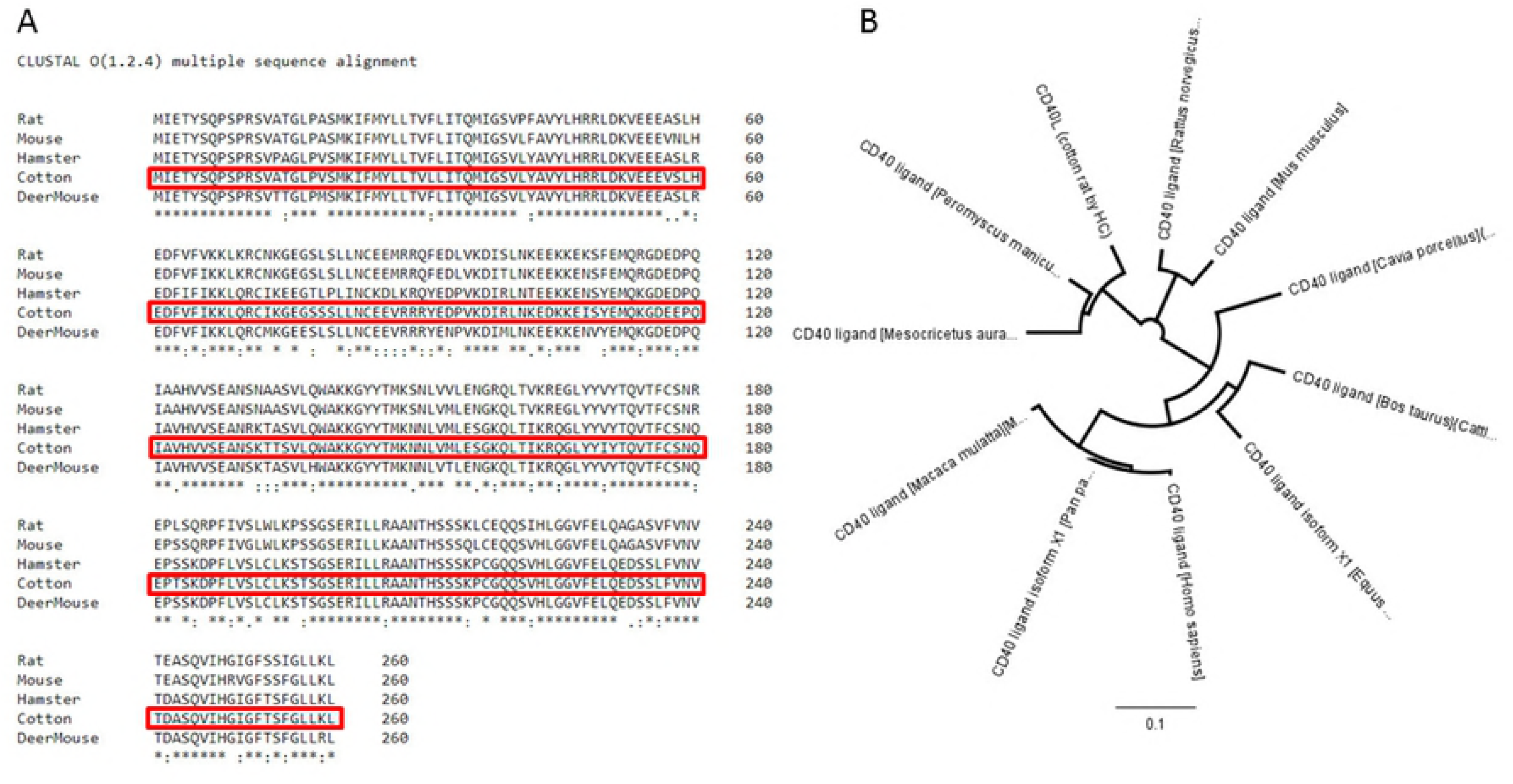
Sequence alignment of Cotton Rat CD40L. (A) The Clustal Omega sequence alignment program from EMBL-EBI was used to align the protein sequence of crCD40L with those of other closely related species. Rat (*Rattus norvegicus* UniProt: Q9Z2V2), Mouse (*Mus musculus* UniProt: P27548), Golden Hamster (*Mesocricetus auratus* XM_005084522.3), and Deer Mouse *Peromyscus maniculatus bairdii* (Accession: XP_006992033). An * (asterisk) indicates positions which have a single, fully conserved residue. A: (colon) indicates conservation between groups of strongly similar properties - scoring > 0.5 in the Gonnet PAM 250 matrix. A. (period) indicates conservation between groups of weakly similar properties - scoring =< 0.5 in the Gonnet PAM 250 matrix. (B) Alignment tree was produced using Geneosis software and multiple alignment was conducted with phylogenetic analysis.

We next examined the functional domains in crCD40L in comparison with other known CD40L. As shown in Fig 3a, crCD40L has a putative tumor necrosis factor (TNF) superfamily domain (aa 137-260) and a 23 aa putative transmembrane domain (aa 23-45). Amino acids 124, 169, 171, 223, 228, 254, and 258 comprise the putative trimer interface, while amino acids 140, 141, 146, 189, 196, and 200 comprise the putative receptor binding sites.

**Fig 3:**
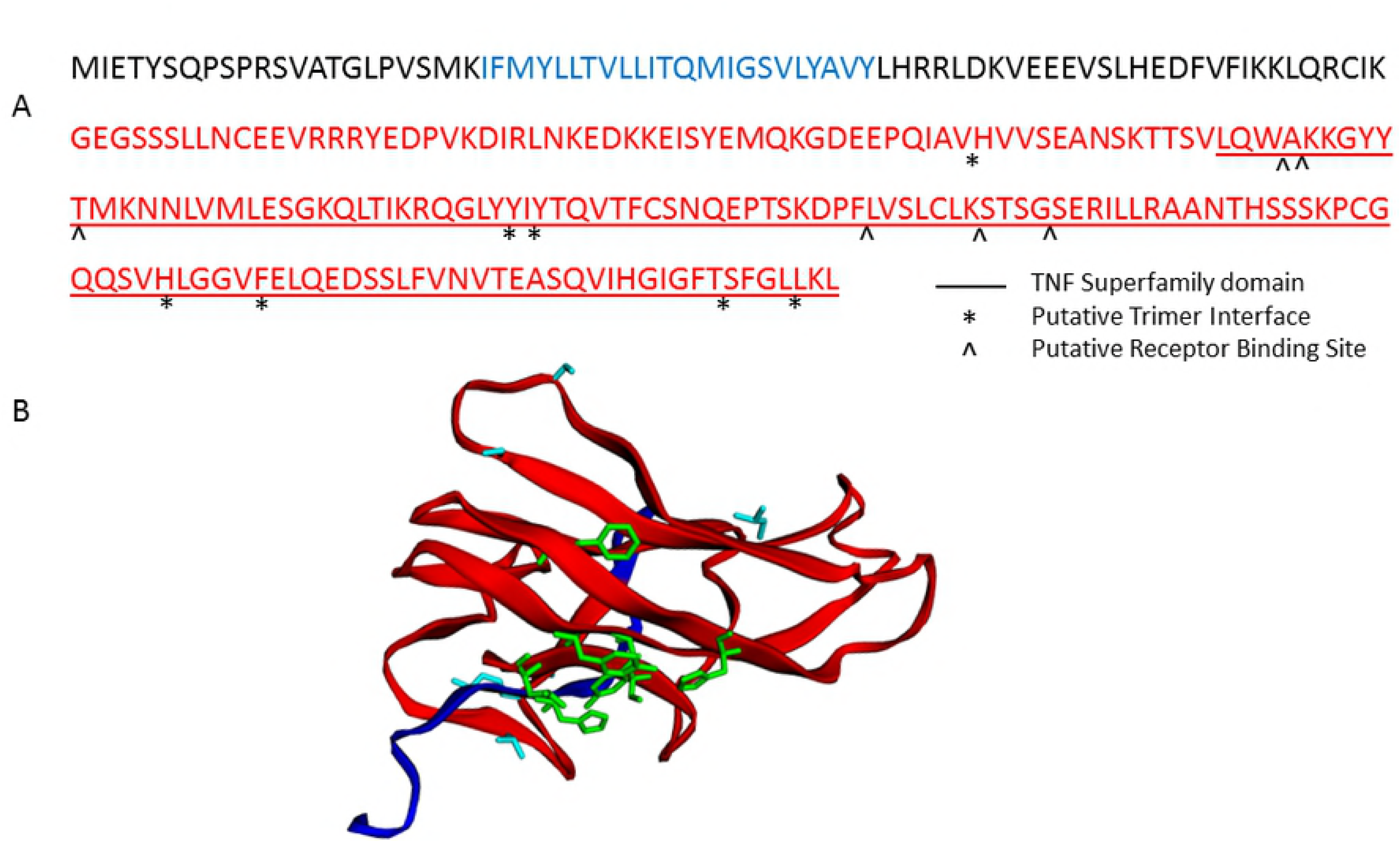
Cotton Rat CD40L putative conserved domains. (A) (-) A line below the sequence indicates the putative TNF superfamily domain. (*) Putative trimer interface on conserved domain TNF. (^) Putative receptor binding sites on conserved domain TNF. The putative transmembrane domain is shown in blue. The putative ecto domain from aa 115 through to aa 260 is shown in red. (B) Image of monomer of Cotton Rat CD40L putative ectodomain conserved regions. The putative TNF superfamily domain is shown in red. (*) Putative trimer interface on conserved TNF superfamily domain residues are shown in green. (^) Putative receptor binding sites on conserved TNF superfamily domain are shown in light blue.

Using EZmol software [35], we predicted folding of the protein as shown in Fig 3b.The cotton rat CD40L cDNA that we have isolated was a 1104 nucleotide sequence with a poly-A tail containing an ORF of 783bp which coded for a 260 aa protein. The homology of cotton rat CD40L, at both the amino acid and nucleic acid level, is closer to members of the Cricetidae family (hamster and deer mouse) than to those of the Muridae family (rat and mouse) as shown in Fig 2b. As with other known CD40L proteins, there is a putative TNF superfamily domain, a transmembrane domain, trimerization sites, and receptor binding sites [36].

TNF superfamily members include TNF (TNF-alpha), LT (lymphotoxin-alpha, TNF-beta), CD40 ligand, Apo2L (TRAIL), Fas ligand, and osteoprotegerin (OPG) ligand, among others [37]. The TNF superfamily is composed of 19 ligands and 29 receptors, in which each has vastly diversified roles in the body and exhibit pro-inflammatory activity, partly via activation of NF-kB [37]. Members of this family generally have an intracellular N-terminal domain, a short transmembrane segment, an extracellular stalk, and a globular TNF-like extracellular domain of about 150 residues [23]. They initiate apoptosis by binding to related receptors, some of which have intracellular death domains [38]. These proteins typically form homo- or hetero-trimeric complexes and bind one elongated receptor molecule along each of three clefts formed by neighboring monomers of the trimer and ligand trimerization is for receptor binding [23] [39] [40]. All seven known conserved residues that constitute the trimer interface on the conserved TNF domain [23] [40] were mapped to the putative crCD40L protein sequence. Additionally, all six known conserved receptor binding sites on the conserved TNF domain [23] [40] were mapped to the crCD40L protein sequence.

### 3.2 Expression of recombinant cotton rat CD40L in Vaccinia Virus

In order to further evaluate the crCD40L deduced sequence, the full 783bp ORF of the crCD40L was cloned into a vaccinia virus vector. The crCD40L construct was designed to carry a secretion signal, histidine tag, and a trimerization motif (Fig 4a). Selection and purification of the recombinant vaccinia virus expressing the CD40L construct was conducted in BHK21 cells. Western blot with anti-histidine antibody (Ab) was used to confirm expression of the CD40L protein construct Fig 4b. The resulting 36 kDa protein product was found in both the cell lysate and supernatant (faint band - 48 hours only). Since the highest expression was found in the cell lysate, it was used for further purification of the protein. It should be noted that the protein was only able to be detected under reducing conditions. Under non-reducing conditions, the protein was unable to be detected by the anti-histidine Ab, even in the cell lysate (data not shown). This indicates that the histidine tag is folded within the trimer and is unavailable in the native form for purification. This is an additional reason for the need to purify the protein from the cell lysate under harsh denaturing conditions followed by protein refolding. The reason we utilized a mammalian expression system to produce the protein rather than a bacterial system is to facilitate its proper folding into its native structure, trimerization, and glycosylation. The aa backbone predicts a protein of 29 kDa, yet initial studies of the CD40L protein suggested a molecular mass of 39 kDa, and on most cell types the molecular mass of CD40L is 32-33kDa, consistent with extensive post-translation modification [36].

**Fig 4:**
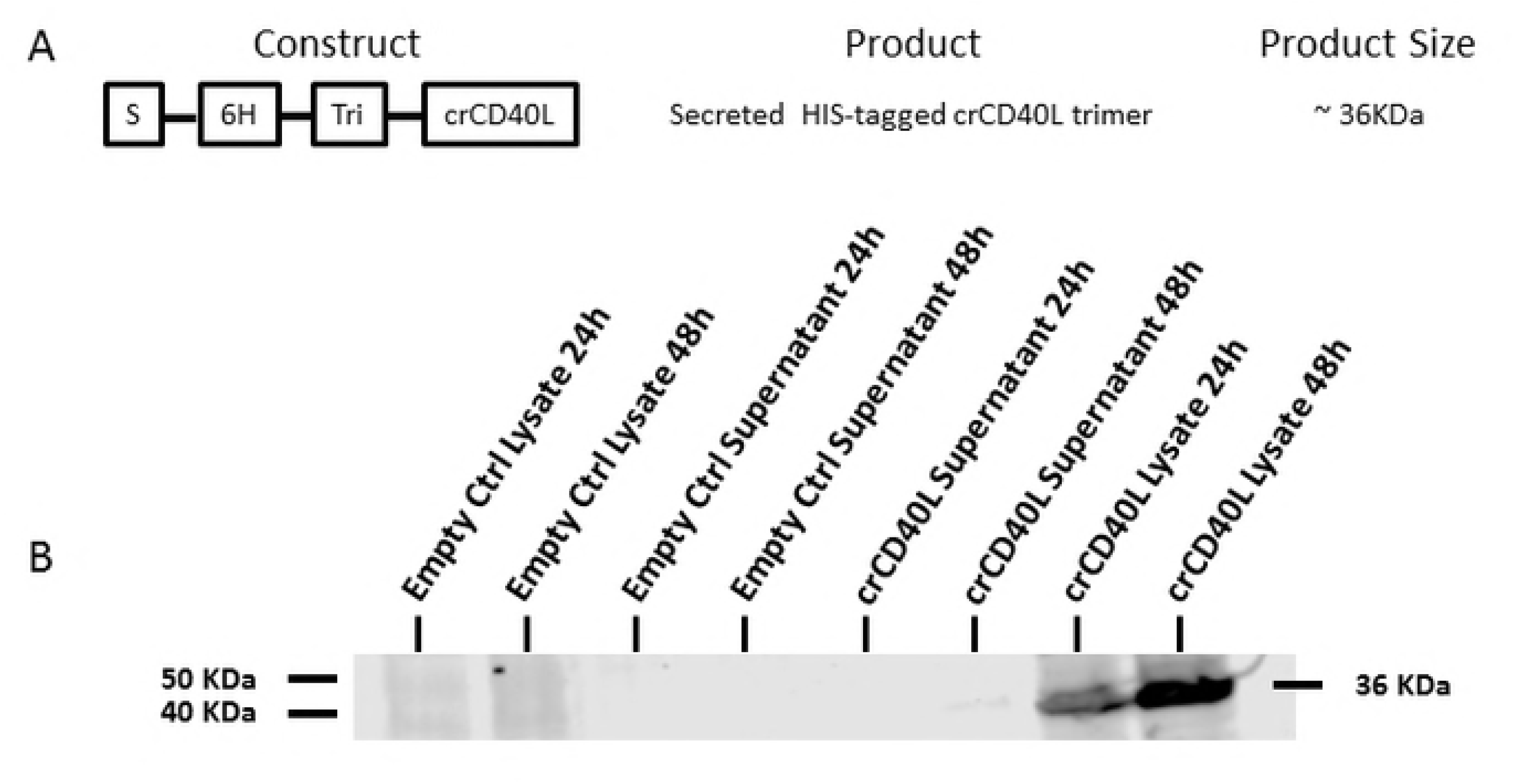
crCD40L construct and protein expression and secretion. (A) Schematic representation of the vv-crCD40L construct. Where “S” is the secretion signal, “6H” is a six histidine residue, and “Tri” is the trimerization motif. (B) In vitro protein expression in BHK21 cell lysate and supernatant collected 24h and 48h post infection. Protein expression is confirmed by Western blot using an anti-histidine Ab.

### 3.3 Purification and verification of cotton rat CD40L

The BHK21 cells expressing the crCD40L construct were collected and lysed with 6 M guanidine hydrochloride with reduced glutathione and sonication. The lysate was bound to a nickel resin and the protein was refolded on the column. Since CD40L biological activity is dependent on a homo-trimer configuration [23], the protein was refolded by gradient, on column buffer exchanges, of a 6M guanidine hydrochloride and phosphate buffer to facilitate gradual removal of the denaturing agent. The resulting bound protein was subsequently eluted with imidazole. The resulting fractions that showed a peak were pooled and dialysed against PBS.

The purified protein was confirmed in ELISA. Since the cotton rat CD40L protein sequence shared 82% identity with the mouse CD40L protein sequence, an Ab known to detect mouse CD40L was used to identify the purified crCD40L protein. The purified recombinant crCD40L was used as a coating antigen, and was detected with an Ab generated against the mouse CD40L Fig 5. Uncoated controls were performed in parallel and were negative for CD40L in ELISA.

**Fig 5:**
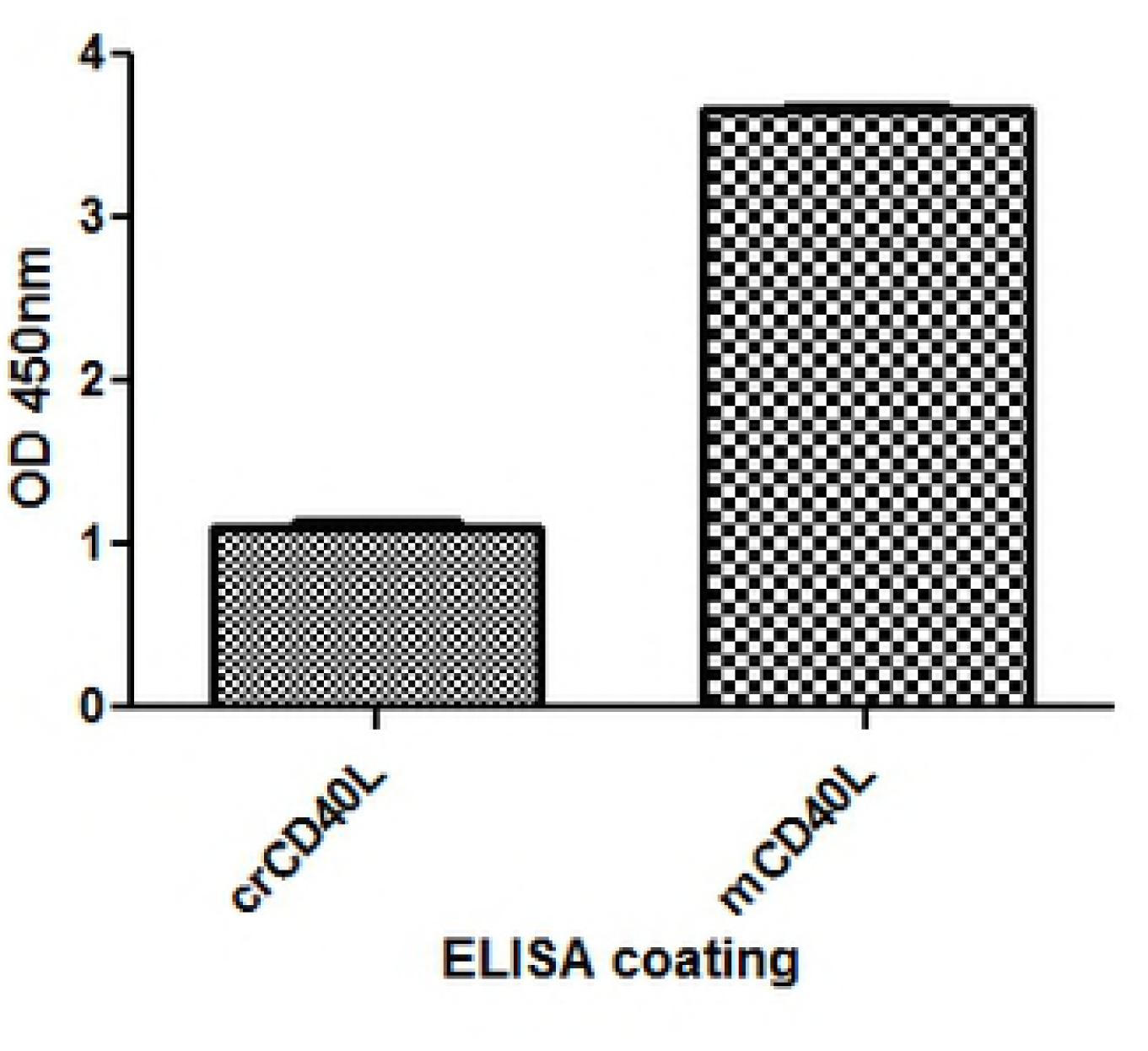
crCD40L is detected by mouse CD40L. crCD40L was expressed in vaccinia virus and purified from infected BHK21 cell lysate on a nickel column. The purified protein was detected by ELISA using a mouse Ab against CD40L.

### 3.4 Functional activity of the recombinant crCD40L

Since the cotton rat CD40L protein sequence shared 82% identity with the mouse CD40L protein sequence with similar functional domains, we evaluated the biological activity of the recombinant crCD40L on immature murine bone marrow DCs. We conducted experiments based on known functional activities of CD40L in other animal species. Specifically, maturation of immature DCs after exposure to antigen is known to play a crucial role in their immunity-stimulating function [36], while trimeric recombinant CD40L has been shown to stimulate DC immunomodulating functions [41]. When CD40L engages CD40 on the surface of DCs, it promotes cytokine production, the induction of cell surface co-stimulatory molecules, and facilitates the cross-presentation of antigen by these cells [27]. In addition, CD11c is a DC integrin marker and upon stimulation, is down-regulated [42]. Intracellular adhesion marker CD54, along with co-stimulatory markers CD40, CD80, and CD86 are all upregulated upon stimulation with CD40L [43] [44]. Moreover, mouse I-A^d^ major histocompatibility complex is also up-regulated upon stimulation with CD40L [44]. When our recombinant crCD40L was used to stimulate immature murine bone marrow DCs, we observed similar results to that when murine CD40L is used (Table1). CD11c was down regulated in both median flouresence intensity and the percentage of positive cells. The co-stimulatory molecules CD54, CD40, CD80, and CD86 were all up-regulated in both median fluorescence intensity and the percentage of positive cells. The Mouse I-A^d^ major histocompatibility complex was upregulated in median fluorescence intensity but not up-regulated in terms of the overall percentage of positive cells. We speculate this to be due to the species incompatibility since we are stimulating mouse bone marrow cells with cotton rat CD40L. Nevertheless, the crCD40L was able to promote up-regulation of key co-stimulatory markers on immature DCs promoting DC maturation.

**Table 1:** crCD40L induces expression of mouse DC maturation markers.

| Marker | Median Fluorescence Intensity |  |  |  | Percentage Positive Cells |  |  |  |
| --- | --- | --- | --- | --- | --- | --- | --- | --- |
|  | U/T | mCD40L<br>3ug/ml | crCD40L<br>3ug/ml | crCD40L<br>30ug/ml | U/T | mCD40L<br>3ug/ml | crCD40L<br>3ug/ml | crCD40L<br>30ug/ml |
| CD11c | 12,131 | 9,921 | 11,515 | 8,960 | 28 | 13 | 23 | 9 |
| CD54 | 2,742 | 2,944 | 3,052 | 2,997 | 0 | 4 | 0 | 2 |
| CD40 | 347 | 794 | 296 | 544 | 57 | 78 | 52 | 69 |
| CD80 | 754 | 1,021 | 743 | 939 | 59 | 71 | 58 | 71 |
| CD86 | 308 | 1,409 | 260 | 748 | 36 | 68 | 34 | 66 |
| I-A<sup>d</sup> | 2,516 | 3,797 | 2,255 | 2,970 | 94 | 95 | 93 | 93 |

U/T: Unstimulated; data are a representative of three separate experiments

CD40-induced activation of cytokine gene expression in DCs by CD40L is an important process in the initiation of primary immune responses and is critical for DC maturation and the generation of antigen-specific T cell responses [45]. IL-6 is a highly pleiotropic cytokine in that it stimulates the activation, proliferation, and survival of T cells, and furthermore, modifies DC function and survival [46] [47] [48] [49]. We tested if the recombinant crCD40L could induce IL-6 gene expression (Fig 6a) and production of the cytokine (Fig 6b) by immature murine bone marrow DCs. The results indicate that a significant increase in both IL-6 gene expression and cytokine production in immature murine bone marrow DCs was observed forty hours after stimulation with the crCD40L. Collectively, the observation that both the upregulation of immature DC cell surface maturation markers and increased IL-6 gene expression and cytokine production provide strong evidence of the biological activity of crCD40L.

**Fig 6:**
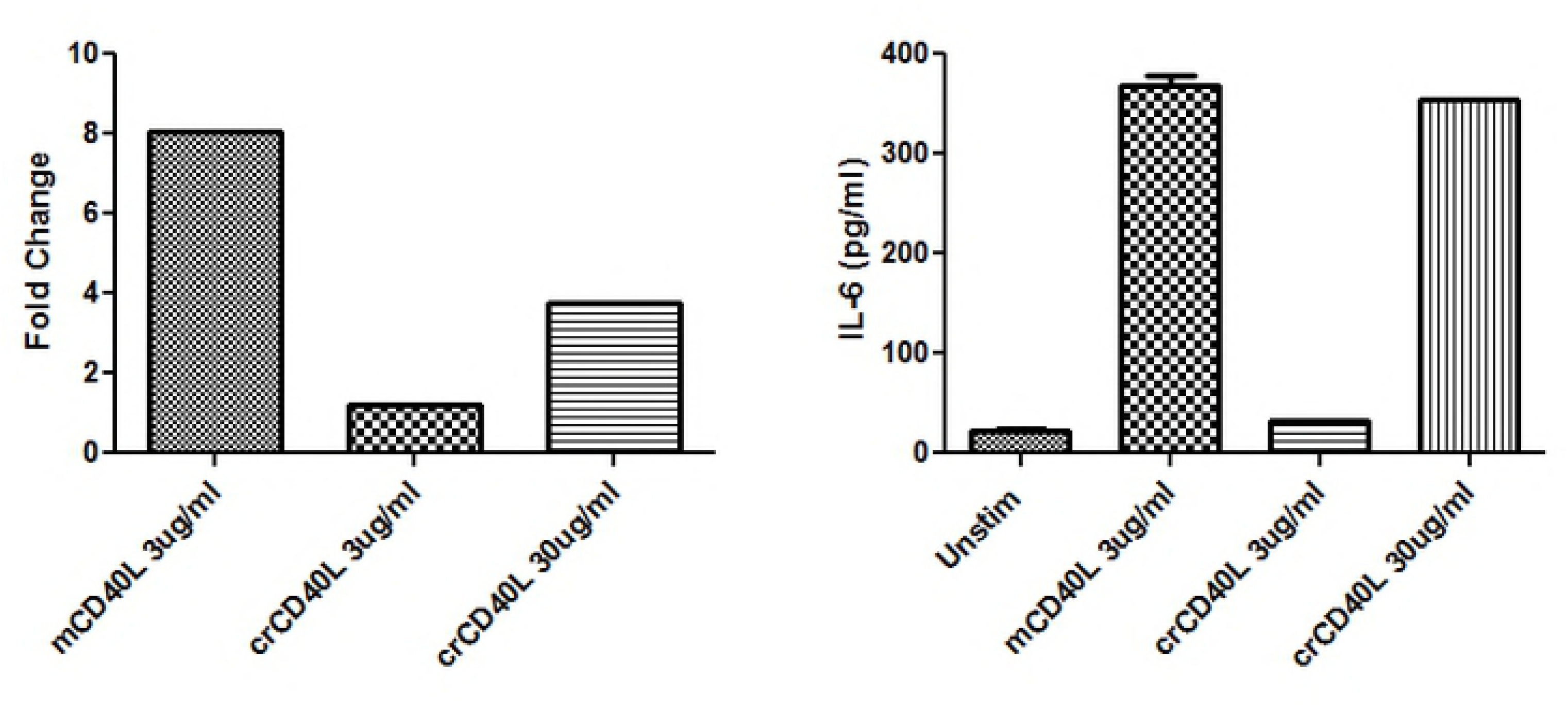
crCD40L induces IL-6 production in immature mouse dendritic cells. Immature mouse dendritic cells were stimulated with either recombinant mouse CD40L, or purified recombinant crCD40L for forty hours. (A) IL-6 gene expression was evaluated using quantitative real-time PCR using 18S to normalize the data. Data are presented as normalized fold change values over unstimulated DCs. (B) IL-6 cytokine levels in the corresponding supernatant were evaluated using ELISA (n=2). These data are a representative of three separate experiments.

In summary, the cotton rat CD40L cDNA that we isolated was a 1104 nucleotide sequence with a poly-A tail containing an ORF of 783 bp which coded for a 260 aa protein. The recombinant cotton rat CD40L was recognized by an Ab against mouse CD40L in direct ELISA, and showed biological activity by upregulating maturation markers (CD40, CD54, CD80, and CD86) as well as I-A^d^ on immature bone marrow murine DCs and moreover, inducing upregulation of IL-6 gene and cytokine expression in these cells.

The isolation of the cotton rat CD40L sequence and availability of CD40L has the potential to positively impact basic immunological research and vaccine development, given the critical importance of this protein in orchestrating immune responses [50] [51].

## Acknowledgementxs

We would like to thank Michele Lemieux, Derek Hodgson for technical assistance.

## References

[1] C. Armstrong, “The Experimental Transmission of Poliomyelitis to the Eastern Cotton Rat, Sigmodon hispidus hispidus,” Public Health Reports, vol. 54, no. 38, pp. 1719–1721, 1939.

[2] R. Faith, C. Montgomery, W. Durfee, E. Aguilar-Cordova and P. Wyde, “The cotton rat in biomedical research,” Lab Anim. Sci., vol. 47, p. 337–345, 1997.

[3] S. Niewiesk and G. Prince, “Diversifying animal models: the use of hispid cotton rats (Sigmodon hispidus) in infectious diseases,” Lab Anim, vol. 36, no. 4, pp. 357–372, 2002.

[4] M. Green, D. Huey and S. Niewiesk, “The cotton rat (Sigmodon hispidus) as an animal model for respiratory tract infections with human pathogens,” Lab Anim (NY), vol. 42, no. 5, pp. 170–176, 2013.

[5] P. R. Wyde, K. J. Stittelaar, A. D. Osterhaus, E. Guzman and B. E. Gilbert, “Use of cotton rats for preclinical evaluation of measles vaccines,” Vaccine, vol. 19, no. 1, pp. 42–53, 2000.

[6] M. Boukhvalova, J. McKay, A. Mbaye, H. Sanford-Crane, J. C. Blanco, A. Huber and B. C. Herold, “Efficacy of the Herpes Simplex Virus 2 (HSV-2) Glycoprotein D/AS04 Vaccine against Genital HSV-2 and HSV-1 Infection and Disease in the Cotton Rat Sigmodon hispidus Model,” J Virol, vol. 89, no. 19, pp. 9825–9840, 2015.

[7] M. G. Ottolini, J. C. Blanco, M. C. Eichelberger, D. D. Porter, L. Pletneva, J. Y. Richardson and G. A. Prince, “The cotton rat provides a useful small-animal model for the study of influenza virus pathogenesis,” J Gen Virol., vol. 86, no. 10, pp. 2823–2830, 2005.

[8] M. C. Eichelberger, “The cotton rat as a model to study influenza pathogenesis and immunity,” Viral Immunol., vol. 20, no. 2, pp. 243–249, 2007.

[9] R. J. Langley, G. A. Prince and H. S. Ginsberg, “HIV type-1 infection of the cotton rat (Sigmodon fulviventer and S. hispidus),” Proc Natl Acad Sci U S A., vol. 95, no. 24, p. 14355–14360, 1998.

[10] M. S. Boukhvalova and J. C. Blanco, “The cotton rat Sigmodon hispidus model of respiratory syncytial virus infection,” Curr Top Microbiol Immunol., vol. 372, pp. 347–358, 2013.

[11] G. A. Prince, D. D. Porter, A. B. Jenson, R. L. Horswood, R. M. Chanock and H. S. Ginsberg, “Pathogenesis of adenovirus type 5 pneumonia in cotton rats (Sigmodon hispidus),” J Virol., vol. 67, no. 1, pp. 101–111, 1993.

[12] J. C. Tsai, G. Garlinghouse, P. J. McDonnell and M. D. Trousdale, “An experimental animal model of adenovirus-induced ocular disease. The cotton rat.,” Arch Ophthalmol., vol. 110, no. 8, pp. 1167– 1170, 1992.

[13] D. D. Porter, G. A. Prince, V. G. Hemming and H. G. Porter, “Pathogenesis of human parainfluenza virus 3 infection in two species of cotton rats: Sigmodon hispidus develops bronchiolitis, while Sigmodon fulviventer develops interstitial pneumonia,” J. Virol., vol. 65, pp. 103–111, 1991.

[14] J. V. Williams, S. J. Tollefson, J. E. Johnson and J. E. J. Crowe, “The cotton rat (Sigmodon hispidus) is a permissive small animal model of human metapneumovirus infection, pathogenesis, and protective immunity,” J. Virol., vol. 79, p. 10944–10951, 2005.

[15] M. A. Rosenfeld, W. Siegfried, K. Yoshimura, K. Yoneyama, M. Fukayama, L. E. Stier, P. K. Pääkkö, P. Gilardi, L. D. Stratford-Perricaudet and M. Perricaudet, “Adenovirus-mediated transfer of a recombinant alpha 1-antitrypsin gene to the lung epithelium in vivo,” Science, vol. 252, no. 5004, pp. 431–434, 1991.

[16] M. A. Rosenfeld, K. Yoshimura, B. C. Trapnell, K. Yoneyama, E. R. Rosenthal, W. Dalemans, M. Fukayama, J. Bargon, L. E. Stier and L. Stratford-Perricaudet, “In vivo transfer of the human cystic fibrosis transmembrane conductance regulator gene to the airway epithelium,” Cell, vol. 68, no. 1, pp. 143–155, 1992.

[17] M. G. Ottolini, D. D. Porter, V. G. Hemming, M. N. Zimmerman, N. M. Schwab and G. A. Prince, “Effectiveness of RSVIG prophylaxis and therapy of respiratory syncytial virus in an immunosuppressed animal model,” Bone Marrow Transplant, vol. 24, pp. 41–45, 1999.

[18] M. G. Ottolini, S. R. Curtis, A. Mathews, S. R. Ottolini and G. A. Prince, “Palivizumab is highly effective in suppressing respiratory syncytial virus in an immunosuppressed animal model,” Bone Marrow Transplantation, vol. 29, pp. 117–120, 2002.

[19] R. G. Crystal, “Adenovirus: The First Effective In Vivo Gene Delivery Vector,” Hum Gene Ther, vol. 25, no. 1, pp. 3–11, 2014.

[20] X. Sáez-Llorens, E. Castaño, D. Null, J. Steichen, P. J. Sánchez, O. Ramilo, F. H. J. Top and E. Connor, “Safety and pharmacokinetics of an intramuscular humanized monoclonal antibody to respiratory syncytial virus in premature infants and infants with bronchopulmonary dysplasia. The MEDI-493 Study Group,” Pediatr Infect Dis J., vol. 17, no. 9, pp. 787–791, 1998.

[21] R. J. W. C. F. L. S. T. A. S. K. N. C. B. M. M. D. M. A. S. D. G. T. D.-S. C. R. M. Armitage, “Molecular and biological characterization of a murine ligand for CD40,” Nature, vol. 357, no. 6373, pp. 80–82, 1992.

[22] R. J. M. R. D. M. S. I. S. J. A. L. A. A. Noelle, “A 39-kDa protein on activated helper T cells binds CD40 and transduces the signal for cognate activation of B cells.,” Proc. Natl. Acad. Sci. USA, vol. 89, no. 14, pp. 6550–6554, 1992.

[23] M. Karpusas, Y. M. Hsu, J. H. Wang, J. Thompson, S. Lederman, L. Chess and D. Thomas, “2A crystal structure of an extracellular fragment of human CD40 ligand,” Structure, vol. 3, no. 10, pp. 1031–1039, 1995.

[24] L. M. Pinchuk, S. J. Klaus, D. M. Magaletti, G. V. Pinchuk, J. P. Norsen and E. A. Clark, “Functional CD40 ligand expressed by human blood dendritic cells is up-regulated by CD40 ligation,” J Immunol, vol. 157, no. 10, pp. 4363–4370, 1996.

[25] A. C. Grammer, M. C. Bergman, Y. Miura, K. Fujita, L. S. Davis and P. E. Lipsky, “The CD40 ligand expressed by human B cells costimulates B cell responses,” J Immunol, vol. 154, no. 10, pp. 4996– 5010, 1995.

[26] V. Henn, J. R. Slupsky, M. Gräfe, I. Anagnostopoulos, R. Förster, G. Müller-Berghaus and R. A. Kroczek, “CD40 ligand on activated platelets triggers an inflammatory reaction of endothelial cells,” Nature, vol. 391, no. 6667, pp. 591–594, 1998.

[27] S. A. Quezada, L. Z. Jarvinen, E. F. Lind and R. J. Noelle, “CD40/CD154 interactions at the interface of tolerance and immunity,” Annu Rev Immunol., vol. 22, pp. 307–328, 2004.

[28] S. M. F. C. Danese S, “The CD40/CD40L costimulatory pathway in inflammatory bowel disease,” Gut, vol. 53, no. 7, pp. 1035–1043, 2004.

[29] G. J. van Mierlo, A. T. den Boer, J. P. Medema, E. I. van der Voort, M. F. Fransen, R. Offringa, C. J. Melief and R. E. Toes, “CD40 stimulation leads to effective therapy of CD40(-) tumors through induction of strong systemic cytotoxic T lymphocyte immunity,” Proc Natl Acad Sci U S A., vol. 99, no. 8, pp. 5561–5566, 2002.

[30] M. E. Grossmann, M. P. Brown and M. K. Brenner, “Antitumor responses induced by transgenic expression of CD40 ligand,” Hum Gene Ther., vol. 8, no. 16, pp. 1935–1943, 1997.

[31] C. Gravel, C. Elmgren, A. Muralidharan, A. M. Hashem, B. Jaentschke, K. Xu, J. Widdison, K. Arnold, A. Farnsworth, A. Rinfret, G. Van Domselaar, J. Wang, C. Li and X. Li, “Development and applications of universal H7 subtype-specific antibodies for the analysis of influenza H7N9 vaccines,” Vaccine. 2015 Feb 25;33(9):1129-34, vol. 33, no. 9, pp. 1129–1134, 2015.

[32] H. Hauser, L. Shen, Q. L. Gu, S. Krueger and S. Y. Chen, “Secretory heat-shock protein as a dendritic cell-targeting molecule: a new strategy to enhance the potency of genetic vaccines,” Gene Ther, vol. 11, no. 11, pp. 924–932, 2004.

[33] A. M. Hashem, C. Gravel, Z. Chen, Y. Yi, M. Tocchi, B. Jaentschke, X. Fan, C. Li, M. Rosu-Myles, A. Pereboev, R. He, J. Wang and X. Li, “CD40 ligand preferentially modulates immune response and enhances protection against influenza virus,” J Immunol, vol. 193, no. 2, pp. 722–734, 2014.

[34] K. J. Livak and T. D. Schmittgen, “Analysis of relative gene expression data using real-time quantitative PCR and the 2(-Delta Delta C(T)) Method,” Methods, vol. 25, no. 4, pp. 402–408, 2001.

[35] C. R. Reynolds, S. A. Islam and M. J. E. Sternberg, “EzMol: A web server wizard for the rapid visualisation and image production of protein and nucleic acid structures.,” J Mol Biol.

[36] C. van Kooten and J. Banchereau, “CD40-CD40 ligand,” J Leuk Biol, vol. 67, pp. 2–17, 2000.

[37] B. B. Aggarwal, S. C. Gupta and J. Kim, “Historical perspectives on tumor necrosis factor and its superfamily: 25 years later, a golden journey,” Blood, vol. 119, no. 3, p. 651–665, 2012.

[38] P. Fiumara and A. Younes, “CD40 ligand (CD154) and tumour necrosis factor-related apoptosis inducing ligand (Apo-2L) in haematological malignancies,” British Journal of Haematology, vol. 113, pp. 265–274, 2001.

[39] Y. Song and P. Buchwald, “TNF Superfamily Protein–Protein Interactions: Feasibility of Small-Molecule Modulation,” Current Drug Targets, vol. 16, no. 4, pp. 393–408, 2015.

[40] M. C. Peitsch and C. V. Jongeneel, “A 3-D model for the CD40 ligand predicts that it is a compact trimer similar to the tumor necrosis factors,” Int. Immunol, vol. 5, pp. 233–238, 1993.

[41] A. E. Morris, R. L. Remmele Jr., R. Klinke, B. M. Macduff, W. C. Fanslow and R. J. Armitage, “Incorporation of an isoleucine zipper motif enhances the biological activity of soluble CD40L (CD154),” J. Biol. Chem., vol. 274, pp. 418–423, 1999.

[42] H. Singh-Jasuja, A. Thiolat, M. Ribon, M. C. Boissier, N. Bessis, H. G. Rammensee and P. Decker, “The mouse dendritic cell marker CD11c is down-regulated upon cell activation through Toll-like receptor triggering,” Immunobiol, vol. 218, no. 1, pp. 28–39, 2013.

[43] B. J. Masten, J. L. Yates, M. Pollard Koga A and M. F. Lipscomb, “Characterization of accessory molecules in murine lung dendritic cell function: roles for CD80, CD86, CD54, and CD40L,” Am J Respir Cell Mol Biol, vol. 16, pp. 335–342, 1997.

[44] D. Y. Ma and E. A. Clark, “The role of CD40 and CD40L in Dendritic Cells,” Semin Immunol, vol. 21, no. 5, p. 265–272, 2009.

[45] J. Mann, F. Oakley, P. W. M. Johnson and D. A. Mann, “CD40 Induces Interleukin-6 Gene Transcription in Dendritic Cells,” J Biol Chem, vol. 277, no. 19, p. 17125–17138, 2002.

[46] J. Van Snick, “Interleukin-6: An Overview,” Annu Rev Immunol, vol. 8, pp. 253–278, 1990.

[47] S. Akira, T. Taga and T. Kishimoto, “Interleukin-6 in Biology and Medicine,” Adv. Immunol., vol. 54, pp. 1–78, 1993.

[48] J. Banchereau, F. Briere, C. Caux, J. Davoust, S. Lebecque, Y.-J. Liu and K. Palucka, “Immunobiology of Dendritic Cells,” Annu. Rev. Immunol., vol. 18, p. 767–811, 2000.

[49] U. Grohmann, F. Fallarino, R. Bianchi, M. L. Belladonna, C. Vacca, C. Orabona, C. Uyttenhove, M. C. Fioretti and P. Puccetti, “In vivo priming of T cells against cryptic determinants by dendritic cells exposed to interleukin 6 and native antigen,” J Immunol, vol. 167, p. 708–714, 2001.

[50] S. Gurunathan, K. R. Irvine, C. Y. Wu, J. I. Cohen, E. Thomas, C. Prussin, N. P. Restifo and R. A. Seder, “CD40 ligand/trimer DNA enhances both humoral and cellular immune responses and induces protective immunity to infectious and tumor challenge,” J Immunol, vol. 161, p. 4563–4571, 1998.

[51] R. J. Noelle, M. Mackey, T. Foy, J. Buhlmann and C. Burns, “CD40 and its ligand in autoimmunity.,” Ann NY Acad Sci, vol. 815, p. 384–391, 1997.

